# Effect of MisMatch Repair Deficiency on metastasis occurrence modelized in a syngeneic mouse model

**DOI:** 10.1101/2024.11.29.626023

**Authors:** Pierre Laplante, Reginaldo Rosa, Laetitia Nebot-Bral, Jordane Goulas, Sergey Nikolaev, Aymeric Silvin, Patricia L Kannouche

## Abstract

Mismatch repair deficiency leads to high mutation rates and microsatellite instability (MSI-H), associated with immune infiltration and responsiveness to immunotherapies. In early stages, MSI-H tumors generally have a better prognosis and lower metastatic potential than microsatellite-stable (MSS) tumors, especially in colorectal cancer. However, in advanced stages, MSI-H tumors lose this survival advantage for reasons that remain unclear. We developed a syngeneic mouse model of MSI cancer by knocking out the MMR gene *Msh2* in the metastatic 4T1 breast cancer cell line. This model mirrored genomic features of MSI-H cancers and showed reduction in metastatic incidence compared to their MSS counterparts. In MSI-H tumors, we observed an enrichment of immune gene-signatures that negatively correlated with metastasis incidence. Importantly, a hybrid epithelial-mesenchymal signature, related to aggressiveness was detected only in metastatic MSI-H tumors which may explain the worse outcomes after recurrence of MSI tumors compared to MSS. Interestingly, we identified immature myeloid cells at primary and metastatic sites in MSI-H tumor-bearing mice, suggesting that MMR deficiency elicits specific immune responses beyond T-cell activation.

**Significance:** A novel syngeneic mouse model of MSI cancer demonstrates that the immune system regulates MSI cancer cell dissemination, offering an important tool to model advanced stages of human MSI-driven disease.

## Introduction

The DNA mismatch repair (MMR) pathway is a major mechanism that ensures the fidelity of genome replication by recognizing and repairing misincorporated bases during DNA replication. Moreover, the MMR machinery corrects small insertions and deletions (indels) that occur during DNA replication, particularly at repetitive sequences known as microsatellites^1^. MMR deficiency (MMRD) is associated with high microsatellite instability (MSI-H), which is observed in over 15 different tumor types, particularly in colorectal cancer (CRC), endometrial, and gastric cancer^2^. The most common cause of MMR deficiency in sporadic human cancers is somatic hypermethylation of the *MLH1* promoter. In addition, Lynch syndrome-related cases are caused by germline pathogenic mutations in one of the four main MMR genes (*MSH2, MSH6, MLH1*, and *PMS2),* followed by somatic events leading to loss of heterozygosity *(for review, see*^3^). Due to their inability to repair replication-associated errors, MSI-H/MMRD tumors harbor large numbers of single-nucleotide substitutions and frameshift mutations, and are characterized by a high tumor mutational burden (TMB)^4,5^ and increased production of immunogenic frameshift-neoantigens. As a consequence, those tumors often exhibit high infiltration of CD8^+^ cytotoxic T lymphocytes (CTL) and Th1 cells, underpinning their immunogenicity and clinical responsiveness to immune checkpoint blockade (ICB) therapy^6,7^

The prognostic value of the MSI phenotype is highly dependent on the cancer stage^8–10^. The MSI phenotype in early-stage setting is generally associated with a better overall prognosis compared to microsatellite stable (MSS) tumors^8–11^. This improved prognosis has been attributed to the high T-lymphocyte invasion and subsequent anti-tumor immune response triggered by their highly immunogenic neoantigen production^12^. While endometrial cancer rarely metastasize^13^, in metastasis-prone cancers such as gastric and colorectal, the MSI phenotype is associated with lower metastatic potential^14,15^. As exemplified for CRC, the most studied metastatic MSI cancer type, the incidence of MSI-H CRCs is higher among stage II (20%) compared to stage III (12%) and is relatively uncommon among metastatic tumors (4%)^15^. Importantly, several retrospective cohort studies have reported that post-recurrence survival is worse among patients with MSI-H tumors compared to those with MSI-Low/MSS tumors^16–18^. The mechanisms underlying the adverse prognostic implications of MSI-H in recurrent CRCs remain largely unknown and require further biological investigation. Moreover, the recurrence pattern of MSI-H colorectal cancers differs from that of their MSS counterpart. Local recurrences and peritoneal metastases are relatively common among patients with MSI-H CRCs, whereas distant metastasis, usually in the liver and lungs, are the most common sites of MSI-L/MSS CRC recurrence^16,19^.

Our current understanding of the molecular mechanisms underlying the metastatic process in MSI-H tumors is limited. Several metastatic mouse models of colorectal cancer have been generated, but they exhibit a low incidence of metastasis^20^. The development of an orthotopic syngeneic murine model of colorectal cancer using CT26 tumor cells provided an accurate model of CRC with tumors developing in the colorectal microenvironment and metastasizing to the liver^21^. However, this model requires sophisticated and costly surgical procedures, with a low metastasis incidence of approximately 20%.

In this study, we established an orthotopic syngeneic mouse model using the 4T1 cell line, a highly metastatic breast cancer (BC) cell line inactivated for the *Msh2* gene. We hypothesized that, due to the tumor-type agnostic nature of MMR deficiency, our MMRD breast cancer model would recapitulate features of MSI metastatic cancers. We observed that while wild-type tumor cells gave rise to metastases in 100% of mice, MMR-deficient 4T1 cells showed a significant reduction (17.3%) in metastasis development in mice. Interestingly, primary MSI-H tumors with no metastasis exhibited a distinct gene expression profile compared to primary MSI-H tumors that developed metastasis. Moreover, by analyzing immune cell populations from primary and (pre)metastatic sites, we discovered a specific immature myeloid population present only in mice bearing MSI tumors, indicating that the defective DNA mismatch repair pathway triggers a specific immune response beyond T-cell activation.

## Materials and Methods

### Cell line

The mouse 4T1 luciferase-expressing (4T1Luc+) cell line was purchased from the American Type Culture Collection (CRL-2539-LUC2) and were cultured at 37°C under 5% CO2 in RPMI- 640 medium supplemented with 10% fetal calf serum, 100 U.mL–1 penicillin (Gibco) and 100 mg.mL–1 streptomycin (Gibco). *Msh2* was inactivated by CRISPR/Cas9, as previously described^22^. Given that mutations accumulate in each round of DNA replication in MMR-deficient cells, parental 4T1Luc+ cells and 4T1*^Msh^*^2^ *^KO^*Luc+ cells were serially passaged *in vitro* during 6 months prior to transplantation into mice.

### Western blot

For validation of Msh2 inactivation, CRISPR-edited clones were lysed in sodium dodecyl-sulfate (SDS) lysis buffer (50 mM Tris pH 7.5, 20 mM NaCl, 10 mM MgCl2, 0.1% SDS, anti- proteases cOmplete cocktail from Roche) supplemented with 20 U/mL benzonase (Millipore) for 10 min at room temperature. Proteins were quantified with Bradford assay and denatured in Laemmli buffer. Proteins were separated on 8% acrylamide SDS–polyacrylamide gels and transferred on PVDF membranes (Millipore). Membranes were blotted with antibodies directed to the following proteins: MSH2 (rabbit, #Ab70240, Abcam, 1/5000), β-actin (mouse, #A5441, Sigma Aldrich, 1/10,000).

### Mouse studies

Female BALB/c mice aged of 6 weeks were purchased at ENVIGO and were maintained in the animal facility of Gustave Roussy. Experiments were performed in accordance with French government and institutional guidelines and regulations. Inoculation of 50,000 4T1 tumor cells were done in the mammary fat pad (left, n°4) of BALB/c mice. Tumors were measured once a week and tumor volume was calculated as follows: length×width². In addition, 15 µg/mL solution of Luciferin (D-Luciferin, Firefly, potassium salt #119222) was injected intraperitoneally, at a rate of 10 µL/g of body weight, for weekly in vivo imaging in an In Vivo Imaging System (IVIS Spectrum CT, Perkin-Elmer). At least one full body and one thoracic picture were acquired each week for each mouse to monitor primary tumor expression of luciferase and metastasis development, respectively. To assess the presence or absence of metastasis in functional regions of the body, 3D CT scans were acquired, and we regionalized the mouse body as follows: the lung (where foci were observed inside the thoracic cage and above the line of the last rib), the bone (where foci overlapped the skeleton), the brain (where foci were observed inside the cranium, non-overlapping the skeleton), the head and neck region (where foci were observed above the shoulder line, non-overlapping the skeleton), the thoracic region (where foci were observed below the shoulder line, and above the last rib, outside the thoracic cage), the abdominal left region (where foci were observed below the thoracic cage, on the left relative to the spine), the abdominal right region (where foci were observed on the contralateral from the abdominal right), and the lower right (where foci were observed below the abdominal right region). The lower left part of the animal, where the primary tumor was located, was not considered. Representative images of the regions are found in supplementary 1B. Mice were euthanized when limit points were reached according to the French and European laws and regulations for the use of mice for scientific purposes.

### Public databases

The Cancer Genomic Atlas (TCGA)^23^ and Memorial Sloan Kettering - Metastatic Events and Tropisms (MSK-MET)^24^ databases were used to calculate metastasis occurrence per patient and per organs respectively, in MSI CRC patients. Both datasets were downloaded through cBioportal^25^. For the analysis of the MSK-MET database, we only considered patients associated with at least one primary tumor sample.

### Nucleic acids extraction

DNA and RNA were co-extracted from primary tumors by a combination of TRIzol-LS (Thermo Fisher, Reference: 10296010) and AllPrep DNA/RNA Mini Kit (Qiagen, Cat. No. 80204) based extraction. Briefly, tumors were submerged in proprietary 1% β-mercaptoethanol RLT buffer (from AllPrep DNA/RNA Mini Kit), dissociated with the homogenizer POLYTRON System PT 3100 D (Kinematica, 441-0750), equipped with “Standard Dispersing Aggregate” tips (Kinematica, PT-DA 07 / 2EC-F101) for 10 seconds, and immediately immerse in ice. This step was repeated until a homogeneous suspension of cells was observed. The lysate was put through an AllPrep DNA spin column, stored at +4C° and further processed following the manufacturer’s instructions. TRIzol-LS/Chloroform was added to the flowthrough (containing the RNA) and the RNA subsequently precipitated. The DNA was quantified by Qubit (Thermo Fisher, Reference: Q32851), and purity was assessed by spectrometry. The RNA was quantified by Qubit (Thermo Fisher, Reference: Q33221), and quality was assessed by calculating the RNA Integrity Number (RIN) through a 2100 Bioanalyzer (Agilent). Only samples with RIN above seven were kept for RNA sequencing.

### Exome analysis

The exomes were sequenced according to the manufacturer protocols (BGI Tech Solutions, Hong Kong) using the Agilent SureSelectXT Mouse All Exon Kit for library construction. We mapped reads using BWA-MEM (v2.2.1) software to the mm9 mouse reference genome and then used the standard GATK best practice pipeline to process the samples and call somatic mutations. PCR duplicates were removed, and base quality scores were recalibrated using MarkDuplicates and BaseRecalibrator tools (both from GATK v4.0.9.0), respectively. Somatic SNVs and INDELs were called and filtered using Mutect2 (from GATK v4.0.9.0, tumor-only mode), FilterMutectCalls and FilterByOrientationBias (both from GATK v4.0.9.0). Quality control of FASTQ files and mapping was done with FastQC (v0.11.8), Samtools (v1.9) and HSMetrics (from GATK v4.0.9.0). Somatic mutations with PASS flag from Mutect2 were additionally filtered to be supported by at least one read from each strand and at least three reads in total. Mutations with a Variant Allele Frequency (VAF) < 0.05 were removed from downstream analyses. Germline variants were defined as those present in more than eight samples out of 12 and excluded from further analysis. Additionally, all variants with a VAF > 0.75 were also excluded. The contribution of the COSMIC mutational signatures was assessed using the MutationalPatterns (v3.12.0) R package. The MSIscore was calculated using msisensor-pro (v1.2.0, tumor-only mode). To calculate TMB, the total number of somatic non-synonymous mutations was normalized to the total number of sequenced megabases.

### Transcriptomic analysis

#### Pre-processing

Paired-end RNA sequencing was performed according to the manufacturer protocols (BGI Tech solutions, Hong Kong) using Optimal Dual-mode mRNA Library Prep Kit (Yeasen, Cat no. 12301ES98) for library construction. The output FASTQ files were pre-processed using Fastp (v0.23.2), and aligned to mm10 reference genome using STAR (v2.7.1a). The resulting BAM files were indexed with Samtools (v1.9). Reads were quantified per features with HTSeq-counts (v2.0.2) using the reference GTF file from Ensembl version 102. FASTQ and BAM files quality were assessed using FastQC (v0.11.8) and Samtools respectively. Using the EdgeR (v4.0.16) R package, read counts were filtered for low-expressed genes and normalized by library size using the edgeR Trimmed of M (TMM) means method. Principal coordinate analysis (PCoA) was performed with EdgeR plotMDS function, using the top 4000 pairwise most variable genes. Log2 counts per millions (logCPM) were calculated using EdgeR cpm function.

#### Unsupervised clustering

After selection of the top 4000 most variable genes by Mean Absolute Deviation (MAD) of logCPM and Z-score transformed, sample-wise unsupervised clustering was performed using 1-Pearson correlation distance and WardD2 linkage method with the complexHeatmap (v2.18.0) R package. Gene-wise clustering was done using the Partitioning Around Medoid (PAM) algorithm implemented in the cluster (v2.1.6) R package.

#### Over-representation analysis

Over-representation analysis (ORA) was performed on the webserver WebGestalt (https://www.webgestalt.org/), using the authors pre-filled options, on Log2 CPM expression tables. The cancer Meta Program (MP) signatures published by Gavish et al., 2023^26^ were downloaded from the authors supplementary data. Automatic mouse-to-human annotation was performed using MGI Jax informatics Human to Mouse Homology database (http://www.informatics.jax.org/downloads/reports/HOM_MouseHumanSequence.rpt) implemented in a custom R script. The remaining unannotated genes were annotated manually using the MGI database.

#### Gene set enrichment analysis

Gene Set Enrichment Analysis (GSEA) was performed on the Broad Institute GenePattern webserver, with the GSEA module. Briefly, CPM expression datasets were converted to .gct files using Broad Institute Morpheus webserver, and phenotype labels files (.cls) (https://software.broadinstitute.org/morpheus/) were generated. Expression datasets, phenotype labels, and signature files were uploaded to the server, and the analysis ran with pre-filled options, except for permutation types set to “gene_set,” as per the authors’ recommendation for analysis with groups of less than seven samples. The Mouse-ortholog hallmark gene set used for this analysis was downloaded from the MSigDB database (https://www.gsea-msigdb.org/gsea/msigdb/mouse/collections.jsp).

#### Immune repertoire

Immune repertoire analysis was performed using the TRUST4^27^ algorithm, implemented in the RNA-seq tumor Immune Analysis (RIMA) pipeline^28^. All TCR metrics shown here are explained in details in the pipeline description (https://liulab-dfci.github.io/RIMA/). All plots were generated in R version 4.3.2, either with package-specific commands, ggplot2 (v3.5.0) or complexHeatmap.

### Neoantigens predictions

Putative neoantigens binding to MHC-I were predicted using the pVACtools (v4.0.6) pipeline^29^, combining WXS and RNAseq files. In addition to the filtering described in the authors documentation (https://pvactools.readthedocs.io/en/latest/index.html), neoantigens were also filtered on anchoring mutation^30^. Briefly, if an anchoring site was mutated, and the affinity of the mutated antigen was weaker than the corresponding wild-type antigen, the neoantigen was discarded.

### Preparation of immune cell suspensions and analysis

Tumor, blood, bone marrow and lungs were harvested. Tissues were cut into small pieces and incubated in 3mL RPMI containing 10% fetal bovine serum, collagenase type IV (0.2 mg/mL, working activity of 770 U/mg) and DNAse (30 mg/mL) for 30min at 37°C. Digested tissues were then passed through a blunt 19G needle to obtain a homogeneous cell suspension and filtered through a cell strainer of 70μM. The cells were washed with FACS buffer (PBS+ 5% BSA + 2mM EDTA) and centrifuged for 4 min at 600g and 4°C before being resuspended. For the lung and tumor samples, 10^7^ cells were resuspended in 80uL of FACS buffer containing 20μL of anti-mouse CD45 microbeads and incubated for 15 min at 4°C. Next, 5mL of FACS buffer was added, and cells were centrifuged at 600g for 4 min at 4°C. Cells were finally resuspended in 2mL of FACS buffer, and CD45+ cells were collected using the Automacs “possel” function followed by centrifugation at 600g for 4 min at 4°C. Cells were resuspended in 200uL of FACS buffer and were mixed at a 1:1 ratio with Whole Blood Cell Stabilizer (Cytodelics), incubated at room temperature for 10 min, and transferred to −80°C freezer to await analysis. These samples were secondarily thawed in a water bath set to 37°C. Cells were fixed at a ratio 1:1 with Fixation Buffer (Cytodelics, ratio 1:1) and incubated for 10 min at room temperature. Red blood cells were lysed by adding of 2mL of Lysis Buffer (Cytodelics, ratio 1:4) at room temperature for 10 min. Cells were washed with 2mL of Wash Buffer (Cytodelics, ratio 1:5). Cells were resuspended in 100μL extra-cellular antibody cocktail and incubated at 4°C for 20 min. Antibodies were diluted in blocking buffer (FACS buffer + 1% rat serum + 1% mouse serum). All antibodies were used at 1/200 dilution and applied at 100μL per cell suspension. Cells were incubated for 15 min with antibodies, at 4°C and then washed with 1 mL of FACS buffer, centrifuged at 600g, for 4 min at 4°C and resuspended in 200μL of FACS buffer. All antibodies are listed in the Supplementary table 2. Samples were acquired on CyTEK Aurora flow cytometer (Cytek Biosciences). Fcs files were exported and analyzed using FlowJo software. Marker expression values were transformed using the auto-logicle transformation function from the flowCore R package. Uniform Manifold Approximation and Projection (UMAP) were carried out using all markers (flow cytometry) or significant PCs (based on Seurat analysis for scRNaseq data). UMAP was run using 15 nearest neighbors (nn), a min_dist of 0.001 to 0.2 and Euclidean distance (Becht et al., 2019; McInnes et al., 2018). Clusters were annotated by the detection of commonly used cell markers, each detailed in Supplementary 3A.

### Statistical analysis

Statistical analyses were performed using functions from the rstatix (v0.7.2) R package. Specifically, Fisher’s exact test of proportions was utilized to determine the statistical significance of metastasis occurrences per animal, patient, and site via the fisher_test() function. For TCR metrics involving comparisons between two groups, the Wilcoxon rank-sum test was applied using wilcox_test(), while comparisons among three groups employed the Kruskal-Wallis test via kruskal_test(). To assess TANs and TAMs percentage population changes between different groups, the Kruskal-Wallis test followed by Dunn’s post-hoc test was conducted using the dunn_test() function. Over-representation analysis and GSEA statistical significance were assessed using WebGestalt (https://www.webgestalt.org/WG_2024_manual.pdf) and GenePattern (https://www.gsea- msigdb.org/gsea/doc/GSEAUserGuideFrame.html) tools, respectively; detailed descriptions of their statistical methodologies are available in their user manuals.

## Results

### Establishment of a MMRD metastatic mouse model

To track metastasis and better understand the mechanisms underlying cancer cell metastasis in the context of MSI phenotype, *Msh2* gene was inactivated using CRISPR/Cas9 in the 4T1 breast cancer cell line expressing luciferase, and Msh2 protein level was assessed by Western Blot (Supplementary 1A). 4T1-*Msh2*KOluc+ cells underwent successive passages in culture over six months, allowing them to accumulate somatic mutations in their genome and indels in microsatellite sequences, as previously described^22^. Both the 4T1-Msh2KO (4T1-MSI) and 4T1-WT (4T1-MSS) cells were orthotopically injected into the lower left mammary fat pad of mice (Figure 1A), and metastasis was monitored using *in vivo* bioluminescence imaging with a luciferase reporter. Twenty-two mice were injected with 4T1-MSS cells, and sixty mice with 4T1-MSI cells. However, 15 out of the 60 4T1-MSI injected mice were excluded from the analysis, mainly due to spontaneous tumor regression. Therefore, only 45 mice bearing 4T1- MSI tumors were included in this study (Figure 1B).

**Figure 1:**
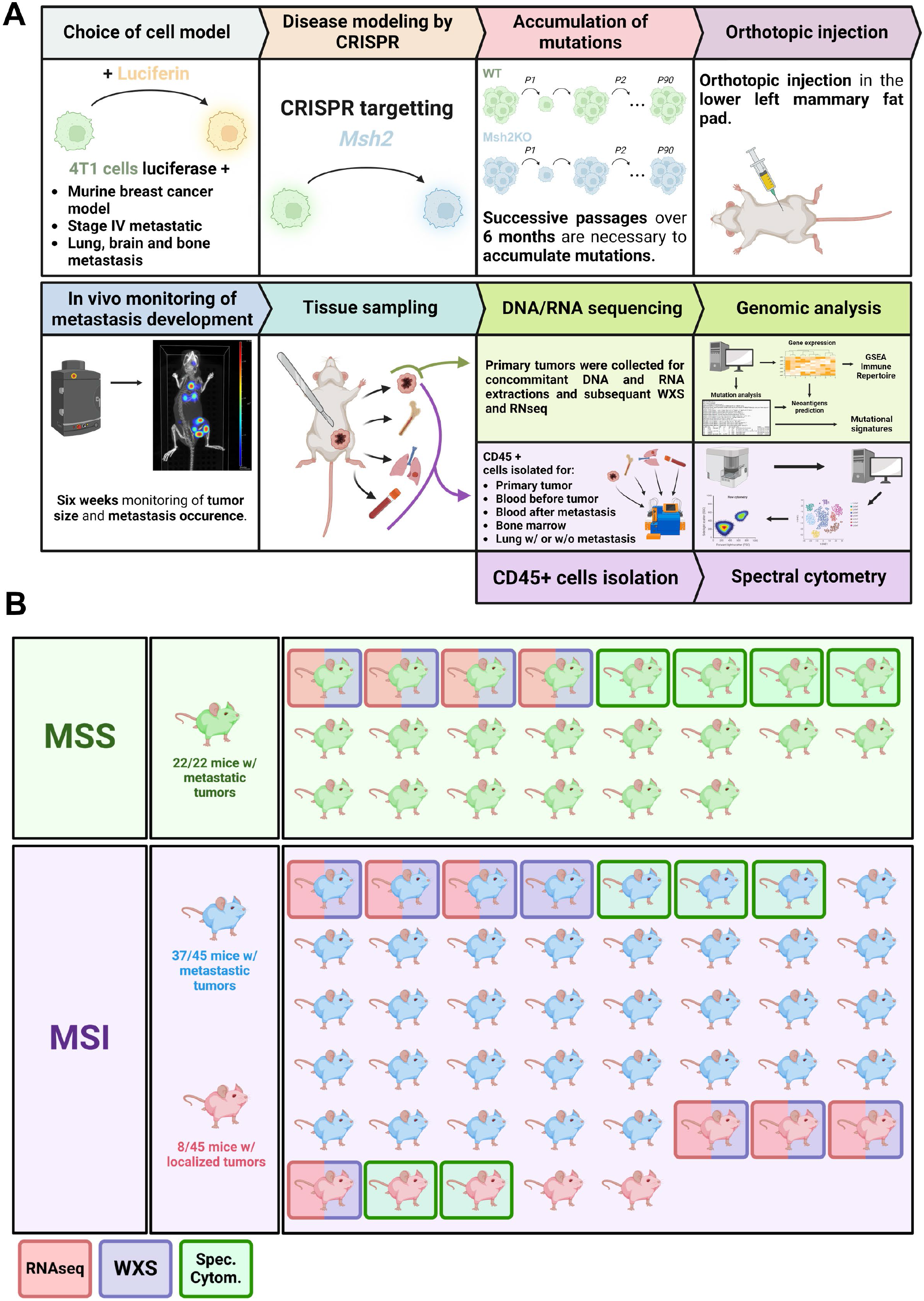
Overall experimental design of the study **A.** Luciferase-expressing 4T1 cell line was inactivated for *Msh2* gene by the CRISPR/Cas9 approach to create MMRD/MSI-4T1 cells. To mimic late-stage MSI cancer, MMRD (and WT, parental 4T1 cells) cells were cultivated for six months *in vitro* to accumulate replication errors. Next, 50,000 WT and MMRD tumor cells were injected into the lower left mammary fat pad of Balb/c female mice. From 1 to 6 weeks post-injection, tumor size was monitored for each mouse, and metastasis incidence was assessed through In Vivo Imaging of luciferase-luciferin bioluminescence (IVIS) before mice sacrifice and tissue harvesting. **B.** Schematic representation of the mice cohort: 82 mice were injected with tumor cells, including 22 mice with WT-4T1 cells and 60 mice with MMRD-4T1 cells. Out of the 22 mice with WT-4T1, 22 developed metastases (green mice). Out of the 60 mice with MMRD-4T1 cells, 15 were removed for technical reasons (see results section). Out of the 45 MSI tumor-injected mice remaining, 37 developed metastasis (blue mice) and 8 did not (red mice). Mice with red squares and/or blue squares were used in the RNA and DNA sequencing respectively. Mice with green square were used in the spectral cytometry experiment. MSS: Microsatellite stable; MSI: Microsatellite instable; WXS: Whole exome sequencing; Spec. Cytom.: Spectral cytometry; w/: with.

### The 4T1-MSI model closely mirrors the genetic and metastatic characteristics of human MSI colorectal cancer

To investigate the metastatic potential of the 4T1-MSI model, we monitored growth of the primary tumor and luciferase signal in injected mice weekly from day 7 to day 50 post-injection of tumor cells. The murine breast cancer model 4T1 is known to primarily metastasize to the lungs. For each mouse, we acquired whole-body and upper-body zoom images to detect the first occurrence of metastasis (Figure 2A). We found that 22 out of 22 (100%) 4T1-MSS animals developed metastases, whereas metastasis was detected in 37 out of 45 (82.3%) 4T1- MSI mice within the same timeframe (Figure 2B). The significant decrease (17.3%, p = 0.0456, Fisher’s exact test) in metastasis occurrence of the 4T1-MSI group highlights the lower ability of MMR-deficient tumors to induce metastasis. We then compared our results to a cohort of colorectal tumors by extracting data from The Cancer Genome Atlas (TCGA) database, we stratified colorectal tumor samples by MSI status and observed a significant discrepancy between early and late-stage cancers. Specifically, 17% of MSS tumors and only 5% of MSI tumors were metastatic (Figure 2B). This finding aligns with published data^15^, further supporting that the MSI phenotype is linked to reduced metastatic potential, as confirmed by our 4T1-MSI model.

**Figure 2:**
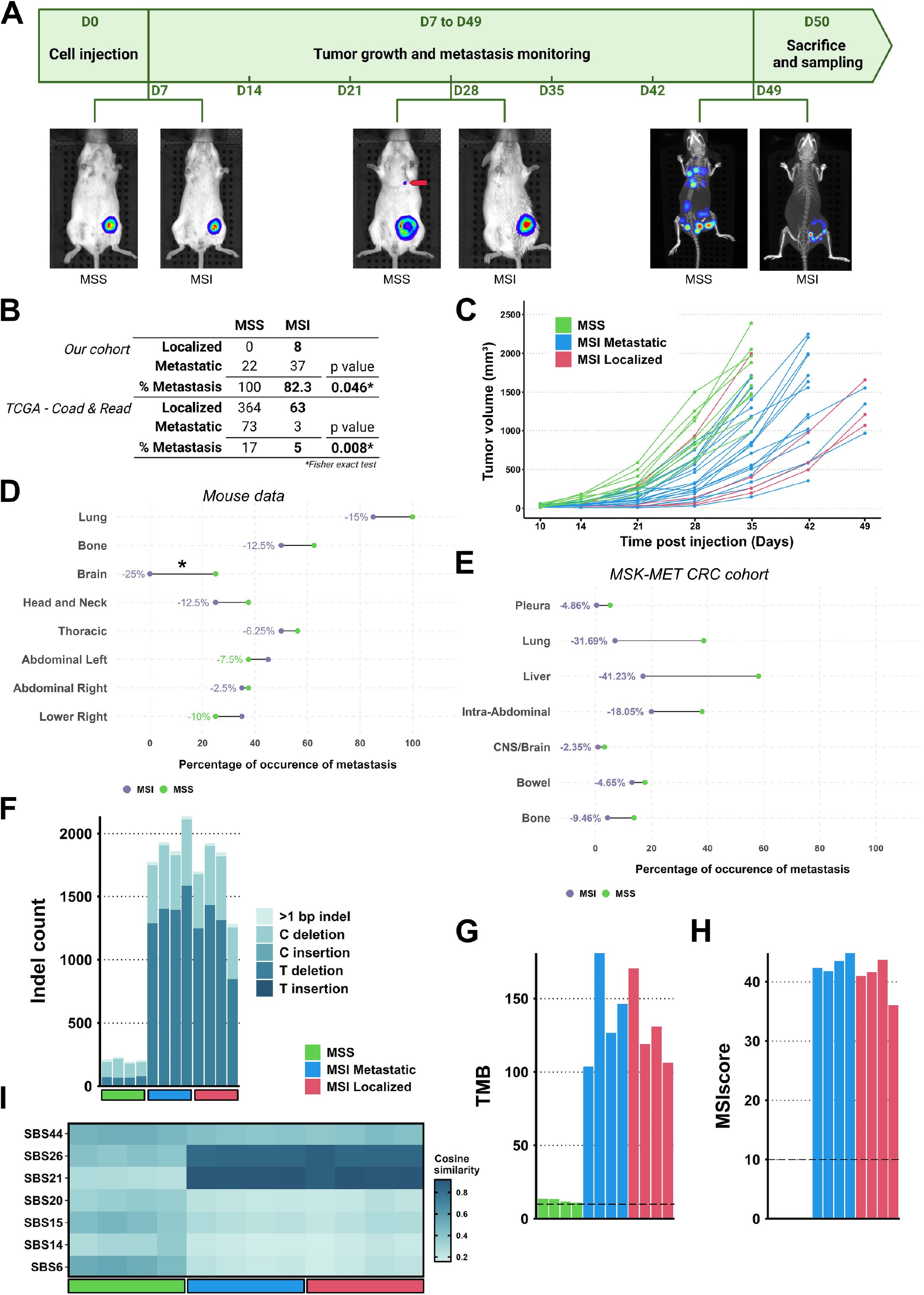
The 4T1-MSI mouse model recapitulated several features of human MSI cancers **A.** 4T1 cells were transplanted into the left mammary fat pad on day 0. Bioluminescence imaging and tumor size measurement was performed at 7 days post-injection and weekly for 6 weeks. Mice were then sacrificed, and tissues were collected. Representative IVIS images of 4T1-WT (MSS, left) and 4T1-MSI tumors (right) are shown at different time points. By day 49, metastasis was only observed in mice with MSS tumors. **B.** Tables displaying the proportion of metastasis occurrences in our cohort and the TCGA COAD/READ cohort, comparing mice (top) and patients (bottom) with or without metastasis. Metastasis detection was done via CT scan, luminescence, and necropsy (for mice), or extracted from TCGA clinical data. P-values were calculated using Fisher’s exact test of proportions. **C.** Line plot showing tumor volume over time (in mm³) for a subset of the mice cohort (n=32). MSS tumors are shown in green, while MSI Metastatic and Localized are shown in blue and red, respectively. **D.** Lollipop plot displaying the percentage of mice with metastasis at each site (described in the methods). The percentage difference in metastasis occurrence between MSS (green) and MSI (purple) is shown, with significance calculated using Fisher’s exact test of proportions. *: p-value < 0.05. **E.** Metastasis burden was transformed into metastasis occurrence (binary data), and the percentage of occurrence per organ was calculated for patients from the MSK-MET CRC database. MSS patients are displayed in green while MSI patients are displayed in purple. All comparisons were significantly different, using Fisher exact test of proportions, p-value < 0.001. **F-H.** Bar plots representing: the count of 1bp and >1bp indels per sample (F), the number of non-synonymous SNVs per Mb (G) and MSIscore (H). The cutoff values for TMB-high (G) and MSI-high (H) were 10 non-syn mut/Mb and 10% of instable microsatellites, respectively (horizontal dotted line). **I.** Heatmap of cosine similarity of the mutational landscape of each sample to MMRD COSMIC signatures. TCGA: The Cancer Genome Atlas; COAD: Colon Adenocarcinoma; READ: Rectal Adenocarcinoma; CT: Computed Tomography; MSK-MET: Memorial Sloan Kettering Metastasis Events and Tropism; Indels: Insertion-deletion; SNVs: Single Nucleotides Variants; Mb: Megabase; TMB: Tumor Mutational Burden.

Based on this observation, the animals were stratified in three distinct groups: 4T1-MSS (referred to as MSS, all metastatic), 4T1-MSI metastatic (referred to as MSI Metastatic), and the 4T1-MSI non-metastatic group (referred to as MSI Localized). Monitoring primary tumor volume over time showed that MSS tumors grow faster than MSI tumors (Figure 2C). However, at equivalent tumor volume, some MSI tumor-bearing mice did not develop metastasis.

To further explore the differences in metastasis occurrence, we divided the mice CT scans into eight functional anatomic regions, as detailed in the methods and Supplementary Figure 1B. Due to the relatively low resolution of the luciferase signal, which limits precise quantification of individual metastatic spots, we opted not to measure the metastasis burden (i.e., the number of metastases per organ). Instead, we focused on quantifying metastasis occurrence at each site, recording only the presence or absence of metastases. We observed that the metastatic MSI group showed a global decrease in metastasis occurrence at the sites of expected tropism (lung, bone, and brain) with a reduction ranging from 12.5% to 25% (Figure 2D). Notably, no brain metastasis was detected in the MSI group, while 25% of MSS mice developed brain metastasis. This result is consistent with the lower rate of metastasis occurrence reported in the literature for MSI versus MSS colorectal cancer^11^. We then compared our findings to the Memorial Sloan Kettering Metastasis (MSK-MET) database, which quantified metastasis occurrence across various sites in a cohort of 2,062 metastatic CRC patients (Figure 2E). When comparing metastasis occurrence between MSS (1832 patients) and MSI-H metastatic CRC patients (230 patients), we observed a reduction in metastasis at all sites in the MSI-H group. This suggests that our murine model of MSI breast cancer faithfully replicates this well- established feature of MSI CRC. Interestingly, when we segregated MSS CRC patient samples by high (157 patients) and low (1,675 patients) TMB (using the FDA-approved immunotherapy biomarker threshold of 10 mutations per megabase^31^), no significant difference in metastasis occurrence was observed (Supplementary Figure 1C). This indicates that the reduced metastasis in MSI tumors is not simply attributable to their high TMB, but rather a specific feature of the MSI phenotype.

While the 4T1-MSI model has been previously well genetically characterized^22^, no data were available on our 4T1-MSI-Luc+ model. Therefore, we performed whole exome sequencing (WXS) on four MSS, four MSI Metastatic, and four MSI Localized primary tumors. Analysis of WXS showed a significantly higher number of indels in the MSI compared to the MSS samples (Figure 2F). We confirmed that the TMB in MSI tumors was elevated compared to MSS samples, ranging between 100 and 175 mutations per megabase (Figure 2G). Consequently, the MSI score, defined as the percentage of unstable microsatellites, was calculated to be around 40 in the MSI group and close to zero in the MSS group (Figure 2H).

Next, we compared the single nucleotide variant (SNV) spectra of our MSI samples to the mismatch repair deficiency (MMRD)-associated COSMIC signatures and found that they closely resembled signatures 21 and 26 in COSMIC v3 (Figure 2I). Importantly, we did not find significant differences in TMB, indel count, or MSI score between the metastatic and localized MSI subgroups. Altogether, these results demonstrate that our 4T1-MSI tumor model recapitulates both the hallmark genetic features and the metastatic behavior of human MSI cancers.

#### Bulk RNA sequencing analysis uncovered distinct transcriptional programs unique to MSS, MSI Metastatic, and MSI Localized samples

To uncover the underlying mechanisms driving the discrepancy in metastasis occurrence between MSS and MSI tumors, we conducted RNA sequencing (RNA-seq) on eleven primary tumor samples, for which we also obtained WXS data. Principal coordinate analysis (PCoA) revealed two outliers - one MSS and one MSI Localized sample (Supplementary Figure 2A). Given the already limited sample size, we excluded these outliers from further analyses to minimize variability and ensure more robust results. We then performed unsupervised clustering on the top 4000 most variable genes between each sample, identified by Mean Absolute Deviation (MAD). Sample-wise clustering revealed three distinct sample clusters corresponding to our groups: MSS, MSI Metastatic, and MSI Localized (Figure 3A, left). After extracting the genes from each gene cluster, we performed over-representation analysis (ORA) with the signatures described by Gavish and colleagues^26^, who identified 41 tumor transcriptomic signatures called MetaPrograms (MP) from single-cell data analysis (Figure 3A, right). We found that Cluster 1 (MSS-specific) was enriched for “EMT-II and Secreted I” MPs, associated with epithelial-mesenchymal transition and epithelial cell secretion, respectively. Cluster 2 (MSI Localized-specific) was uniquely enriched for the “Interferon/MHC-II” MP, which includes MHC-I genes. Cluster 3 (predominantly MSI Metastatic) was enriched for “Cell Cycle – G1/S and Stress” MPs, containing proliferation and general cellular stress-related genes. Furthermore, cluster 4 mixed MSS and MSI Metastatic gene cluster showed enrichment for EMT-I and Hypoxia signatures, the former being a fully mesenchymal program and the latter a hypoxia/glycolysis program. These results indicate that our clustering classification of samples based on their MSI and metastatic status is linked to distinct transcriptomic profiles. Specifically, the stratification reveals EMT-related gene expression in the MSS group, immune-related genes in the MSI-localized group, and proliferation-related genes in the MSI-metastatic group.

**Figure 3:**
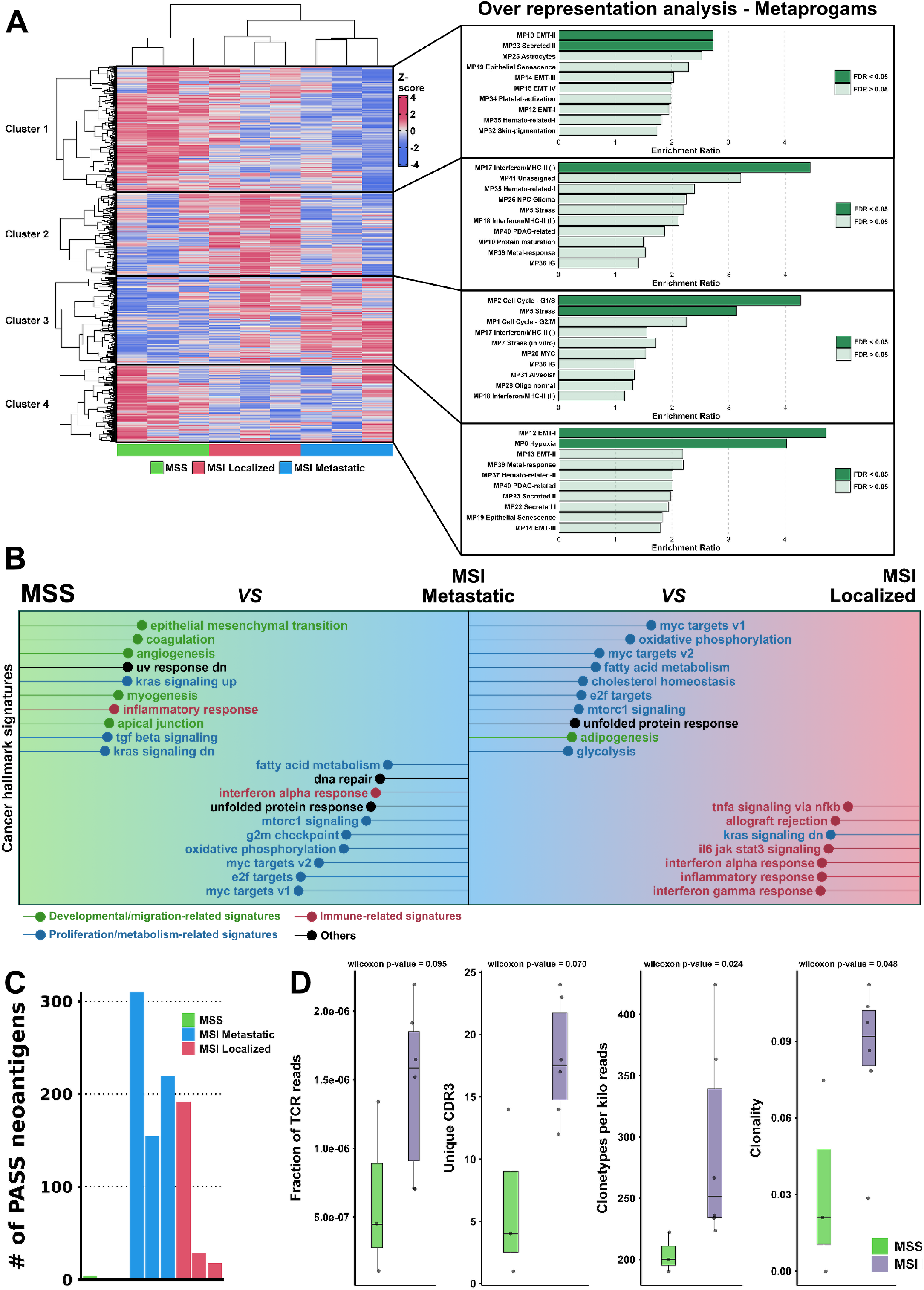
Transcriptomic profiles of MSS, MSI Metastatic, and MSI Localized tumor samples. **A.** Unsupervised clustering of the top 4000 most variable genes (left) and ORA of each cluster for 41 cancer MPs (right). Significant enrichment is represented in dark green. **B.** Pairwise GSEA differential enrichment of MsigDB 50 Hallmark signatures. Left and right part represent two independent comparisons. The length of each line represents the absolute value of normalized enrichment score (NES) obtained for each signature in each comparison. **C.** Barplot of the number of unique putative neoantigens per sample. **D.** Boxplots of TCR repertoire metrics. From left to right: (i) fractions of total reads mapping to the TCR for each sample, (ii) the number of unique CDR3 (hypervariable region of the TCR) per sample, (iii) clonotypes per kilo reads (CPK) representing the number of unique CDR3 divided by the total amount of reads mapped to all CDR3, multiplied by 1000 to obtained normalized number CDR3 per 1000 TCR reads, a marker of diversity and (iv) clonality calculated as 1-Shannon’s entropy/log(N), N being the number of unique CDR3 clones. ORA: Over-Representation Analysis; MPs: Meta Programs; MAD: Mean Absolute Deviation; GSEA: Gene set Enrichment Analysis; TCR: T-Cell Receptor; CDR3: Complementarity-determining region 3.

We then conducted a Gene Set Enrichment Analysis (GSEA) between MSS and MSI tumors using cancer Hallmark signatures. MSS tumors were enriched for “EMT and developmental” signatures, while MSI tumors were enriched for “immune and proliferative” signatures (Supplementary Figure 2B). Based on the results of the unsupervised clustering, we hypothesized that the immune signature enrichment in the MSI group stemmed from MSI Localized samples, and the proliferative signatures from MSI Metastatic samples. Pairwise GSEA confirmed this hypothesis, showing that MSI Localized tumors were enriched for immune signatures relative to MSI Metastatic tumors, while MSI Metastatic tumors were enriched for proliferative signatures compared to MSS tumors (Figure 3B). Surprisingly, EMT signatures were enriched in MSS tumors relative to MSI Metastatic samples but not in MSI Metastatic compared to MSI Localized samples (Figure 3B). Using four different “EMT” MPs (Gavish et al., 2023), we performed pairwise GSEA (Supplementary Figure 2C) and observed that EMT-I (full mesenchymal program per the authors^26^) was enriched in MSS tumors relative to MSI Metastatic tumors, while EMT-III (hybrid epithelial-mesenchymal) was enriched in MSI Metastatic tumors relative to MSI Localized tumors. The hybrid E/M state is associated with increased tumor cell plasticity during the metastasis process^32^. MSS tumors showed enrichment for multiple EMT states, whereas MSI Metastatic tumors were predominantly EMT-III, indicating a more homogenously aggressive state.

The increased immune activity observed in MSI tumors has been proposed to be driven by neoantigen production and presentation via MHC-I. These neoantigens are recognized by T cell receptors (TCRs), triggering an adaptive immune response^6,7^. We predicted neoantigen binding to MHC-I using the pVACtools pipeline^29^, combining WXS and RNAseq data. After stringent filtering, we observed a sharp increase in predicted neoantigens in MSI tumors compared to MSS tumors (Figure 3C). Intriguingly, MSI Localized tumors exhibited much less neoantigens in two out of three samples as compared to MSI Metastatic tumors. To further understand the implication of neoantigens production in intra-tumoral T-cells, we characterized the immune repertoire of T cell receptors (TCR) from our RNAseq data (Figure 3D and Supplementary Figure 2D). We observed an increase in both clonality and diversity (Clonotype per kilo reads, CPK) of T cells present in MSI tumors compared to the MSS group, in line with the neoantigens load increase (Figure 3D). However, by examining MSI Metastatic and Localized separately, MSI Localized tumors showed a tendency for increased clonality and decreased diversity of TCR compared to MSI Metastatic tumors (Supplementary Figure 2E). This may reflect the influence of a few immunogenic neoantigens private to the MSI Localized group (Supplementary Figure 2D), which drove the clonality of intra-tumoral TCR. Altogether, these findings demonstrate that our model replicates key immunological aspects of human MSI tumors, including enhanced immune response markers likely due to increased neoantigen production, and associated with TCR diversity/clonality.

#### Analysis of immune cell populations in the 4T1-MSI tumor microenvironment revealed MSI-specific myeloid populations

To obtain an overview of the immune landscape in our MSI cancer model, we performed spectral cytometry on the CD45+ cells from various compartments of nine mice (four mice transplanted with 4T1-MSS and five with 4T1-MSI, of which three mice developed metastases and two did not). We collected peripheral blood before tumor injection and at endpoint (i.e., at 6 weeks of tumor growth), as well as the primary tumor, lung (with and without metastasis), and bone marrow (BM) samples. Post-sacrifice, tissues were dissociated, and CD45+ cells were enriched using CD45 microbeads and marked with a panel of 26 antibodies targeting different cell surface markers to identify main immune cell types (Figure 4A, details in Methods).

**Figure 4:**
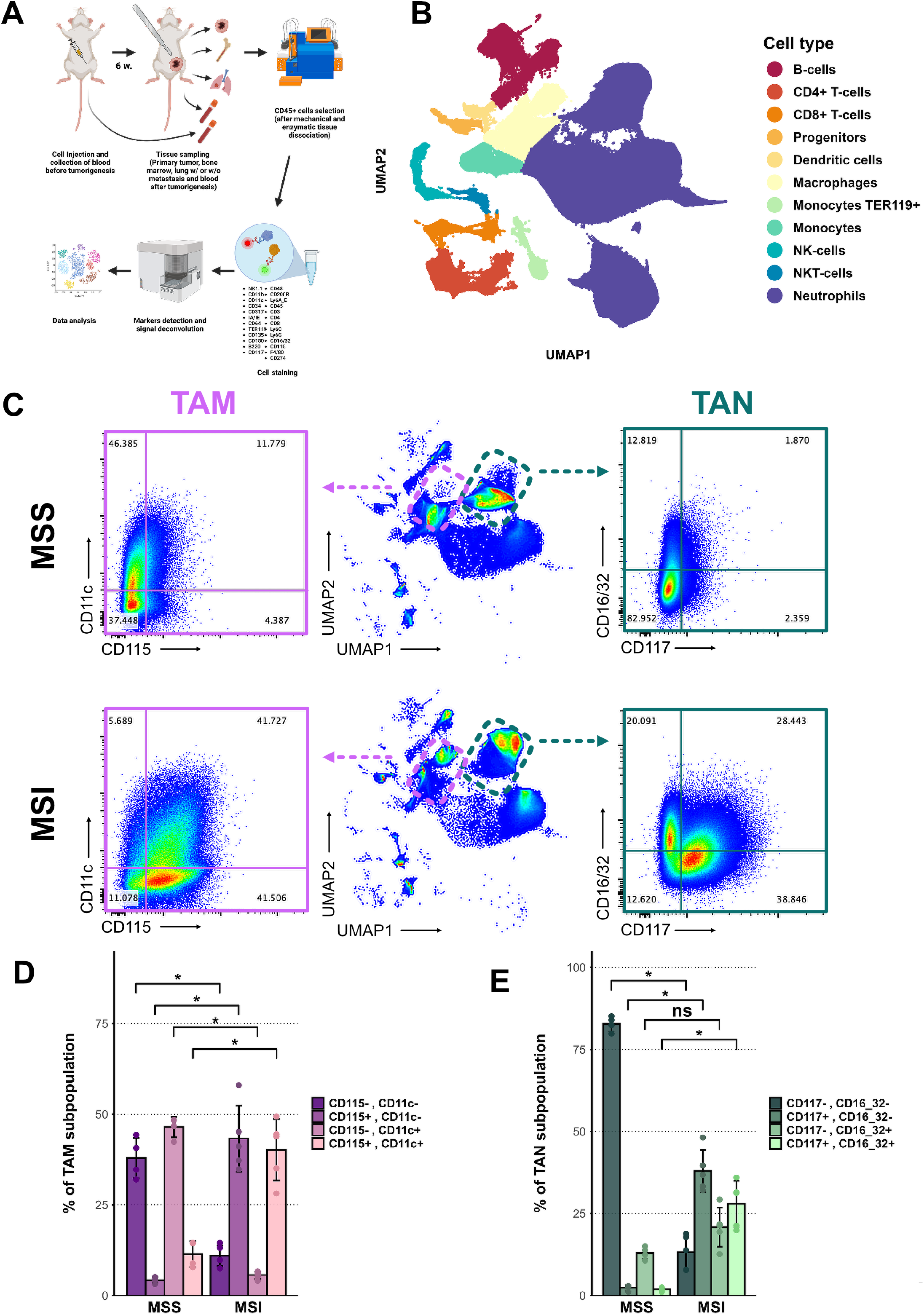
Analysis of immune cell populations in the tumor microenvironment of MSS, MSI Metastatic, and MSI Localized samples. **A.** Experimental design of the spectral cytometry experiment: After injection of 50,000 WT and MMRD-4T1 cells in the left mammary fat pad of the mice, metastases development was monitored for 6 weeks, and the primary tumor, bone marrow, lungs, and blood collected before tumor injection and at endpoint were recovered for CD45+ cell enrichment. Each immune compartment was marked with a panel of antibodies (see supplementary table II). After the detection and deconvolution of the markers, the results were represented through Uniform Manifold Approximation and Projection (UMAP) for dimension reduction. Clusters of cell populations were annotated manually, according to each marker expression, shown in supplementary 3A. **B.** UMAP of total events from all samples in all immune compartments**. C.** FACs plots showing the expression of CD11c and CD115 in TAM (purple) and the expression of CD16/36 and CD117 in TAN (green) in MSS and MSI tumors. **D-E.** Bar graphs reporting the percentages of TAM and TAN subpopulations identified in C. Significance was determined through Wilcoxon rank-sum test, error bars represent standard deviation, *: p-value < 0.05. TAM: Tumor-associated macrophages; TAN: Tumor- associated neutrophils.

We performed Uniform Manifold Approximation and Projection (UMAP) analysis to visualize clustering patterns within each immune compartment (Figure 4B). Quantification of neutrophil clusters showed a significant increase in neutrophil populations across all compartments following tumor injection, highlighting their expansion in response to tumor presence. Indeed, before tumor injection, neutrophils accounted for no more than 0.3% of CD45+ cells per sample in the blood, whereas post-tumorigenesis, neutrophils comprised between 57% and 96% of CD45+ cells across all compartments (Supplementary Figure 3B and supplementary Table 1). This increase is consistent with previous findings on the 4T1 breast cancer model^22,33^. We then investigated the differences in immune cell populations between MSS and MSI samples. While blood and BM samples did not show significant variation in immune cell populations between the two groups (Supplementary Figure 3C), a dramatic shift was observed in tumor-associated neutrophils (TANs) and macrophages (TAMs) within the primary tumor (Figure 4C, D, and E). We observed differential expression of TANs and TAMs surface markers in MSI tumors compared to MSS tumors (Figure 4C). A subpopulation of TANs in MSI tumors showed increased expression of CD117 (cKit), a marker of early myeloid progenitors. These CD117+ TANs were 20 times more prevalent in MSI tumors compared to MSS tumors (Figure 4C right and E). Furthermore, the subpopulation of CD16/32+, CD117- active neutrophils was more prevalent in MSI tumors, consistent with a richer immune activity. Regarding TAMs, a 10-fold increase in CD11c-, CD115+ cells was observed in MSI tumors (Figure 4C left and D, I). CD115, a marker usually expressed on monocytes that binds CSF-1 to promote survival and differentiation into macrophages, is associated with poor prognosis when overexpressed in tumors^34–37^. The increased prevalence of these TAMs appears to be specific to the MSI phenotype. We did not observe significant differences in the overall immune composition between MSI Metastatic and MSI Localized samples, likely due to the small sample size and high variability within the MSI Localized group. Nevertheless, these findings demonstrate that our model effectively enables the study of the tumor microenvironment (TME) in MSI cancer, enabling the identification of distinct immune subpopulations associated with the MSI phenotype. Intriguingly, no significant differences were observed in the relative abundance of T cells (CD4+ and CD8+) between MSS and MSI groups at the primary tumor site (Supplementary Figure 3E, left) or at the metastatic site (Supplementary Figure 3E, right). However, RNA-seq analysis revealed increased cytotoxic activity in MSI tumors (Supplementary Figure 3D). These findings suggest that, despite similar T-cell quantities, the gene expression profile in MSI tumors reflects enhanced cytotoxic activity, indicating functional differences in the immune response.

### The 4T1-MSI model enables the identification of immature neutrophils in lungs free of metastasis development

The presence of metastatic and non-metastatic mice transplanted with MSI-4T1 cells allowed for the unique opportunity to characterize the microenvironment of the lung, which is the main metastatic site in our model within an MSI context. Analysis of markers expressed by immune cell populations in lung samples revealed that the previously identified population of MSI-specific TAMs and TANs were identifiable in the lungs of mice from the MSI groups (Figure 5A). Although these immune cell populations were present in very low numbers (ranging from 0.2% to 0.01% of the total lung cell CD45+ populations in the MSI groups), they exhibited the same patterns of marker expression observed in the primary tumors (Figure 5A). Intriguingly, the TAN-MSI subpopulation observed at the primary tumor site was also present in the lungs of both metastatic and non-metastatic mice (Figure 5A right and B). Conversely, the TAM-MSI subpopulation was only detected in the microenvironment of metastatic MSI mice (Figure 5A left and B). These findings suggest in our model reveals the presence of an MSI-specific immature neutrophil population in the primary tumors, which is detected in the lungs, and may exist independently of metastatic development. Altogether, these results indicate that our 4T1- MSI model is well-suited for identifying and characterizing MSI-specific subpopulations of immune cells at metastatic sites.

**Figure 5:**
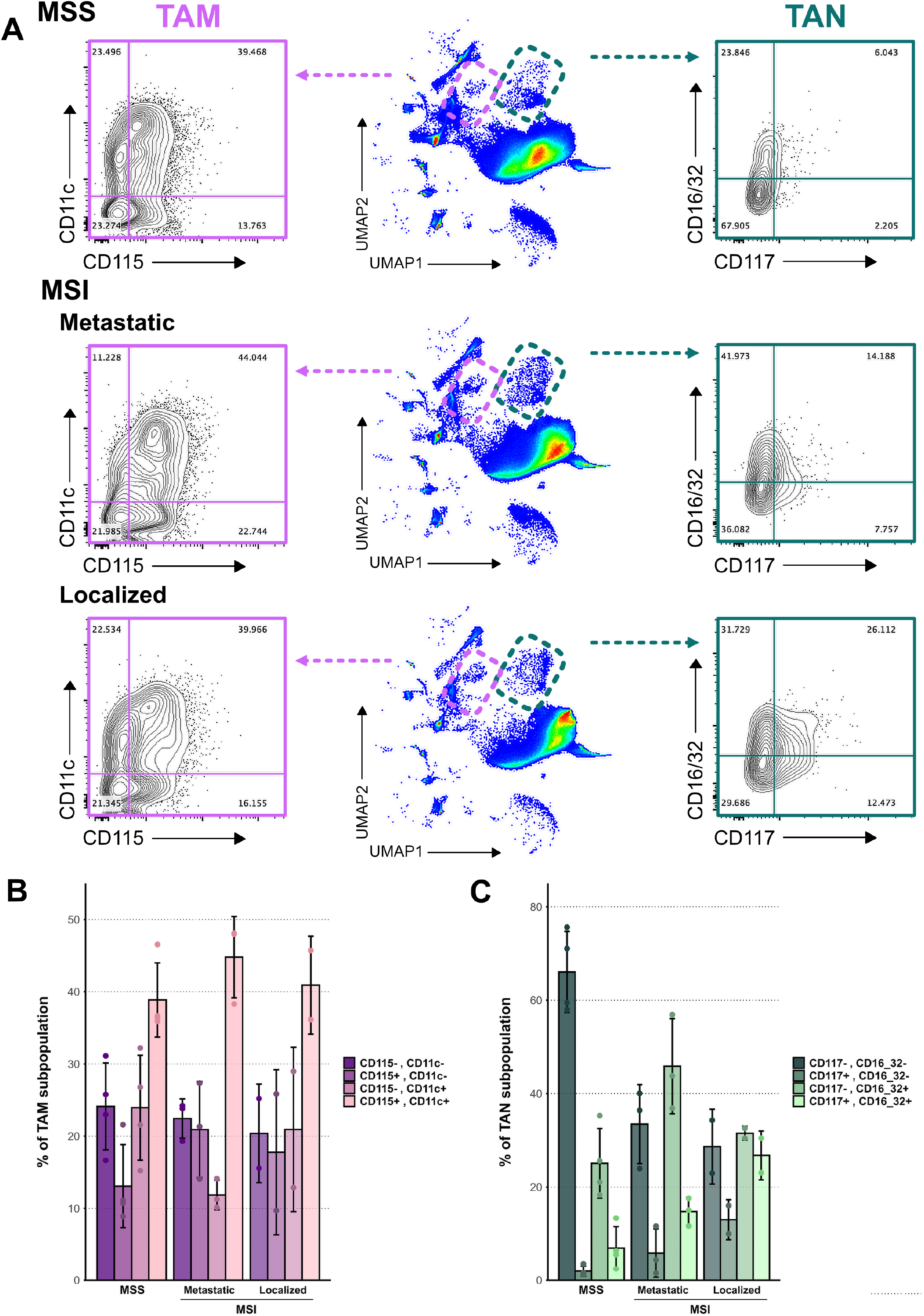
Identification of MSI-specific myeloid populations at the secondary sites of mice bearing MSI tumors. **A.** FACs plots showing the expression of CD11c and CD115 (TAM in purple) and the expression of CD16/36 and CD117 (TAN in green) in the lung of MSS, MSI Metastatic, and MSI Localized samples. From top to bottom, the MSS, MSI Metastatic and MSI Localized lung samples are shown, with TAMs and TANs markers (on the right and left of the graph respectively) defined in Figure 4. The middle is the UMAP of the CD45+ cells from the lungs, for each group. **B-C.** Bar graphs reporting the percentages of TAM (B) and TAN (C) subpopulations identified in A. TAN populations are expressed as percentage of TANs, while TAMs are percentage of CD45+ cells. Error bars represent standard deviation. TAM: Tumor-associated macrophages; TAN: Tumor-associated neutrophils.

## Discussion

In this study, we successfully established a mouse model of MSI metastatic cancer using the orthotopic 4T1 breast cancer cell line with *Msh2* gene inactivation. This MMR-deficient model demonstrated a reduction of metastatic incidence by 17.3% compared to the MMR-proficient model and a reduction in lesion-bearing sites per mouse in the MSI Metastatic group compared to the MSS group. These findings align with human metastatic MSI CRC data from the TCGA and MSK-MET databases. Although our MSI model showed an accumulation of SNVs specific to two MMRD COSMIC signatures, along with increased indels, TMB, and MSI score, these features alone did not explain the differences between metastatic and localized MSI tumors. Instead, metastasis incidence in the MSI context was associated with distinct RNA-seq signature enrichment: immune-related in the MSI Localized group and proliferative in the MSI Metastatic group. Curiously, we did not find enrichment of the classical EMT signature in MSI Metastatic samples compared to MSI Localized samples. Instead, we identified a unique enrichment of a hybrid EM signature, suggesting that MSI tumors may undergo EMT in a distinct manner, compared to MSS tumors. Neoantigen prediction and T-cell receptor (TCR) repertoire analysis revealed increased neoantigen load and greater TCR diversity and clonality in MSI group compared to MSS group. Interestingly, the MSI Localized group showed a lower neoantigen load but a tendency toward higher clonality than the MSI Metastatic group. These findings imply that early selection by cytotoxic CD8+ T-cell activity against tumor cells expressing immunogenic neoantigens resulted in lower neoantigen burden and a restricted TCR repertoire in MSI Localized compared to MSI Metastatic samples. While TCR diversity is typically associated with better outcomes^38^, Riaz and colleagues demonstrated a correlation between TCR clonality and a lower neoantigen burden in melanoma patients who responded to ICB compared to non-responders^39^. A detailed analysis of the tumor microenvironment revealed MSI-specific markers of immaturity expressed by TANs and TAMs. Interestingly, MSI-specific TANs, but not TAMs, were found in the lungs of both metastatic and non-metastatic animals suggesting that immature neutrophils may be present at metastatic sites prior to the development of metastasis. Overall, our findings indicate that MMR deficiency may influence the tumor immune landscape beyond intra-tumoral T-cells.

Our MSI syngeneic mouse model revealed two distinct profiles of MSI tumors: (i) metastatic MSI tumors, characterized by higher proliferative capacity, an aggressive hybrid epithelial-mesenchymal (E-M) state, and lower intra-tumoral immune activity, and (ii) localized MSI tumors, marked by lower proliferative capacity, no EMT activity, and higher intra-tumoral immune activity, particularly associated to interferon γ and α signatures. This heterogeneity mirrors findings by Kim JH and colleagues^40^, who reported significant variability in tumor-infiltrating lymphocyte density and tertiary lymphoid structure in human MSI-H CRC, without notable differences in TMB or neoantigen load. In their study, “immune-high” MSI-H CRC was linked to lower recurrence and metastasis rates, while “immune-low” MSI-H CRC was associated with higher recurrence and metastasis. Our BC model reflects this heterogeneity despite the difference of sites of primary tumors, underscoring the agnostic nature of MMR deficiency on tumor fate. Immune activity at the primary site appears to be a key predictor of recurrence and metastasis in MSI cancers. At the population level, the MSI-H/MMRD phenotype alone may suffice to predict metastatic potential in CRC. However, at the individual level, assessing intra-tumoral immune activity, including lymphocyte infiltration and immune markers, is essential for accurate prognosis. Our findings suggest that attempts to turn "cold" tumors to "hot" by inhibiting the MMR pathway may have mixed outcomes, necessitating careful monitoring of tumor-infiltrating immune cells and potential shifts in metastatic sites.

Tumors can promote the abnormal presence of immature myeloid cells outside the bone marrow^41^ by either inducing the premature release of immature cells from the bone marrow^42^ or by limiting the differentiation of monocytes in situ^43^, which is often associated with a poor prognosis. A recent study on intra-tumoral neutrophils in a mouse model of pancreatic cancer demonstrated the presence of both mature (T2) and immature (T1) neutrophils within tumors, which eventually progressed to pro-tumoral neutrophils (T3)^44^. In our study, we identified tumor-associated neutrophils and macrophages that appear specific to MSI tumors and exhibit markers of immaturity. We hypothesize that these neutrophils may represent T1-like neutrophils, which could eventually differentiate into T3-like pro-tumoral neutrophils. Similarly, the immature macrophages observed in our analysis may be in the process of transitioning into pro-tumoral M2 macrophages, which are known to express CD115 and rely on its signaling pathway for polarization towards the M2 phenotype^45^. Further investigations, in other mouse models and human samples, are needed to determine the precise role of these aberrant myeloid cells. Nevertheless, the presence of these immature TANs in the lungs of both metastatic and localized MSI tumor-bearing mice suggests a distinct mechanism by which MSI tumors may establish distant metastatic sites, separate from MSS tumors. Validating these findings with human data would be diagnostically valuable but remains challenging due to the limited availability of metastatic MSI samples.

Our model is not intended to replicate an MSI CRC model but rather to serve as a tool for improving our understanding of MMRD/MSI-H phenotype in a metastatic context. We believe that the agnostic nature of MMR deficiency enables the investigation of the metastatic process within an MSI framework using a BC model, despite the fact that MSI is observed in no more than 2% of human BC cases^46^. The 4T1 cell line is an ideal cancer model for monitoring metastasis development because (i) it is highly metastatic and exhibits a pronounced tropism, (ii) it can be transplanted into immunocompetent mice, and (iii) it can be injected orthotopically into the mammary fat pad of the mice, mimicking the pathology of breast cancer and promoting metastasis development, in contrast to subcutaneous models which are less prone to induce metastases^47^. This 4T1-MSI mouse model remarkably mirrors several key observations relevant to MSI CRC. Indeed, this study highlights the ambivalence of MSI status concerning prognosis: on one hand, we observed reduced aggressiveness in the 4T1 model when microsatellite instability is present (evidenced by slower tumor growth rates, fewer metastases, and higher immune activity); on the other hand, there is a significant enrichment of proliferation and EMT-III (hybrid E-M) signatures in MSI Metastatic samples compared to MSS and MSI Localized samples, respectively. These findings may explain the association of MSI status with a better prognosis compared to MSS, while also offering insight into the worse outcomes of MSI cancers compared to MSS after recurrence^16–18^.

We acknowledge several limitations in this study, including differences in the cell of origin of our model compared to the most prevalent forms of human metastatic cancer. A key characteristic of MSI cancers—the heightened infiltration of T-cells—is not recapitulated in the 4T1-MSI model in this study. Moreover, the flow cytometry panel used was relatively broad, limiting the ability to examine specific immune cell subpopulations in greater detail. Future studies employing single-cell RNA sequencing on primary tumors and metastatic sites in this model could yield invaluable insights into the tumor microenvironment and help identify key players in the metastatic process of MSI tumors. Performing a longitudinal study could also reveal whether the trajectory of immune activity and metastasis in MSI tumors is predetermined or not, and provide insights into potential key factors steering the tumors toward different outcomes. Lastly, the small number of animals used in the genomic and flow cytometry experiments somewhat reduces statistical power. Nevertheless, despite these limitations, we were able to uncover biologically relevant and statistically significant findings regarding MSI metastatic cancers.

Overall, we have characterized a novel syngeneic mouse model of microsatellite instability metastatic cancer, using the 4T1 breast cancer cell line. The data generated have validated the model’s relevance to human MSI metastatic cancers, serving as a foundation for future research on the mechanisms of metastasis in MSI cancer patients.

## Acknowledgments

The authors acknowledge the contribution of Patrick Gonin and Karine Ser-Le Roux from the Preclinical Evaluation PlatForm (PFEP), and Tudor Manoliu from the Imaging and Cytometry PlatForm (PFIC) at Gustave Roussy. The authors express their gratitude to Valerie Rouffiac for her precious help and valuable advice on the IVIS spectrum CT system. We also thank all members of the Kannouche and Nikolaev lab for helpful discussions. This work was performed thanks to Gustave Roussy core facilities.

The P.L.K lab was supported by La Ligue Nationale contre le Cancer (Equipe labellisée). This work was supported by the SIRIC SOCRATE 2.0 INCa-DGOS-INSERM_12551. P.L. received support from the Ministère de l’Enseignement Supérieur et de la Recherche. J.G. received support from la Ligue Nationale Contre le Cancer. L.N.B received funding from Philanthropia Foundation - Lombard Odier and la Ligue Nationale Contre le Cancer. R.R. received support from Campus France «Eiffel Excellence Scholarship Program”.

## Author contributions

R. R. and P.L.K. conceptualized the study, while A.S. designed the experiments for immune cell population analysis. P.L. and R. R. conducted the mouse experiments with the help of J.G.

P.L. performed all additional experiments and bioinformatic analyses, with support from S.N.

L.N.B. initiated the early phase of the project. P.L., A.S. and P.L.K. drafted the manuscript, and all authors reviewed and approved the final version.

## Declaration of Interests

The authors declare no competing interests.

## Data availability

All raw data generated in this study are available upon request from the corresponding author.

## Notes

### Competing Interest Statement

The authors have declared no competing interest.

## References

1. Kunkel, T. A. & Erie, D. A. Eukaryotic Mismatch Repair in Relation to DNA Replication. Annu Rev Genet 49, 291–313 (2015).

2. Dudley, J. C., Lin, M.-T., Le, D. T. & Eshleman, J. R. Microsatellite Instability as a Biomarker for PD-1 Blockade. Clinical Cancer Research 22, 813–820 (2016).

3. Nebot-Bral, L. et al. Hypermutated tumours in the era of immunotherapy: The paradigm of personalised medicine. European Journal of Cancer 84, 290–303 (2017).

4. Alexandrov, L. B. et al. The repertoire of mutational signatures in human cancer. Nature 578, 94–101 (2020).

5. Chalmers, Z. R. et al. Analysis of 100,000 human cancer genomes reveals the landscape of tumor mutational burden. Genome Med 9, 34 (2017).

6. Llosa, N. J. et al. The vigorous immune microenvironment of microsatellite instable colon cancer is balanced by multiple counter-inhibitory checkpoints. Cancer Discov 5, 43–51 (2015).

7. Le, D. T. et al. PD-1 Blockade in Tumors with Mismatch-Repair Deficiency. N Engl J Med 372, 2509–2520 (2015).

8. Wang, B., Li, F., Zhou, X., Ma, Y. & Fu, W. Is microsatellite instability-high really a favorable prognostic factor for advanced colorectal cancer? A meta-analysis. World J Surg Oncol 17, 169 (2019).

9. Guo, Y. et al. The clinicopathological characteristics, prognosis and immune microenvironment mapping in MSI-H/MMR-D endometrial carcinomas. Discov Onc 13, 12 (2022).

10. Beghelli, S. et al. Microsatellite instability in gastric cancer is associated with better prognosis in only stage II cancers. Surgery 139, 347–356 (2006).

11. Han, S. et al. The distinct clinical trajectory, metastatic sites, and immunobiology of microsatellite-instability-high cancers. Front. Genet. 13, (2022).

12. Laghi, L. et al. Prognostic and Predictive Cross-Roads of Microsatellite Instability and Immune Response to Colon Cancer. Int J Mol Sci 21, 9680 (2020).

13. Mao, W. et al. Clinicopathological study of organ metastasis in endometrial cancer. Future Oncology 16, 525–540 (2020).

14. Cristescu, R. et al. Molecular analysis of gastric cancer identifies subtypes associated with distinct clinical outcomes. Nat Med 21, 449–456 (2015).

15. Roth, A. D. et al. Prognostic Role of KRAS and BRAF in Stage II and III Resected Colon Cancer: Results of the Translational Study on the PETACC-3, EORTC 40993, SAKK 60- 00 Trial. JCO 28, 466–474 (2010).

16. Kim, C. G. et al. Effects of microsatellite instability on recurrence patterns and outcomes in colorectal cancers. Br J Cancer 115, 25–33 (2016).

17. Koopman, M. et al. Deficient mismatch repair system in patients with sporadic advanced colorectal cancer. Br J Cancer 100, 266–273 (2009).

18. Tran, B. et al. Impact of BRAF mutation and microsatellite instability on the pattern of metastatic spread and prognosis in metastatic colorectal cancer. Cancer 117, 4623–4632 (2011).

19. Sinicrope, F. A. et al. DNA Mismatch Repair Status and Colon Cancer Recurrence and Survival in Clinical Trials of 5-Fluorouracil-Based Adjuvant Therapy. JNCI: Journal of the National Cancer Institute 103, 863–875 (2011).

20. Romano, G., Chagani, S. & Kwong, L. N. The path to metastatic mouse models of colorectal cancer. Oncogene 37, 2481–2489 (2018).

21. Evans, J. P. et al. Development of an orthotopic syngeneic murine model of colorectal cancer for use in translational research. Lab Anim 53, 598–609 (2019).

22. Nebot-Bral, L. et al. Overcoming resistance to αPD-1 of MMR-deficient tumors with high tumor-induced neutrophils levels by combination of αCTLA-4 and αPD-1 blockers. J Immunother Cancer 10, e005059 (2022).

23. Weinstein, J. N. et al. The Cancer Genome Atlas Pan-Cancer analysis project. Nat Genet 45, 1113–1120 (2013).

24. Nguyen, B. et al. Genomic characterization of metastatic patterns from prospective clinical sequencing of 25,000 patients. Cell 185, 563–575.e11 (2022).

25. Gao, J. et al. Integrative Analysis of Complex Cancer Genomics and Clinical Profiles Using the cBioPortal. Sci Signal 6, pl1 (2013).

26. Gavish, A. et al. Hallmarks of transcriptional intratumour heterogeneity across a thousand tumours. Nature 618, 598–606 (2023).

27. Song, L. et al. TRUST4: immune repertoire reconstruction from bulk and single-cell RNA-seq data. Nat Methods 18, 627–630 (2021).

28. Yang, L. et al. Tutorial: integrative computational analysis of bulk RNA-sequencing data to characterize tumor immunity using RIMA. Nat Protoc 18, 2404–2414 (2023).

29. Hundal, J. et al. pVACtools: a computational toolkit to identify and visualize cancer neoantigens. Cancer Immunol Res 8, 409–420 (2020).

30. Capietto, A.-H. et al. Mutation position is an important determinant for predicting cancer neoantigens. J Exp Med 217, e20190179 (2020).

31. Marcus, L. et al. FDA Approval Summary: Pembrolizumab for the treatment of tumor mutational burden-high solid tumors. Clin Cancer Res 27, 4685–4689 (2021).

32. Liao, T.-T. & Yang, M.-H. Hybrid Epithelial/Mesenchymal State in Cancer Metastasis: Clinical Significance and Regulatory Mechanisms. Cells 9, 623 (2020).

33. Elsas, M. van et al. Host genetics and tumor environment determine the functional impact of neutrophils in mouse tumor models. J Immunother Cancer 8, e000877 (2020).

34. Becker, S., Warren, M. K. & Haskill, S. Colony-stimulating factor-induced monocyte survival and differentiation into macrophages in serum-free cultures. J Immunol 139, 3703–3709 (1987).

35. Ide, H. et al. Expression of colony-stimulating factor 1 receptor during prostate development and prostate cancer progression. Proc Natl Acad Sci U S A 99, 14404– 14409 (2002).

36. Wang, H. et al. Interactions between colon cancer cells and tumor-infiltrated macrophages depending on cancer cell-derived colony stimulating factor 1. Oncoimmunology 5, e1122157 (2016).

37. Chambers, S. K. Role of CSF-1 in progression of epithelial ovarian cancer. Future Oncol 5, 1429–1440 (2009).

38. Bortone, D. S., Woodcock, M. G., Parker, J. S. & Vincent, B. G. Improved T-cell Receptor Diversity Estimates Associate with Survival and Response to Anti–PD-1 Therapy. Cancer Immunology Research 9, 103–112 (2021).

39. Riaz, N. et al. Tumor and Microenvironment Evolution during Immunotherapy with Nivolumab. Cell 171, 934–949.e15 (2017).

40. Kim, J. H. et al. Genomic and transcriptomic characterization of heterogeneous immune subgroups of microsatellite instability-high colorectal cancers. J Immunother Cancer 9, e003414 (2021).

41. Kusmartsev, S. & Gabrilovich, D. I. Immature myeloid cells and cancer-associated immune suppression. Cancer Immunol Immunother 51, 293–298 (2002).

42. Mackey, J. B. G., Coffelt, S. B. & Carlin, L. M. Neutrophil Maturity in Cancer. Front Immunol 10, 1912 (2019).

43. Broz, M. L. & Krummel, M. F. The Emerging Understanding of Myeloid Cells as Partners and Targets in Tumor Rejection. Cancer Immunol Res 3, 313–319 (2015).

44. Ng, M. S. F. et al. Deterministic reprogramming of neutrophils within tumors. Science 383, eadf6493 (2024).

45. Haegel, H. et al. A unique anti-CD115 monoclonal antibody which inhibits osteolysis and skews human monocyte differentiation from M2-polarized macrophages toward dendritic cells. MAbs 5, 736–747 (2013).

46. Bonneville, R. et al. Landscape of Microsatellite Instability Across 39 Cancer Types. JCO Precis Oncol 1, PO.17.00073 (2017).

47. Gómez-Cuadrado, L., Tracey, N., Ma, R., Qian, B. & Brunton, V. G. Mouse models of metastasis: progress and prospects. Dis Model Mech 10, 1061–1074 (2017).

