## Supplemental Figures S1-S3 for "Effect of MisMatch Repair Deficiency on metastasis occurrence modelized in a syngeneic mouse model"

### SUPPLEMENTAL INFORMATION

#### Supplemental Figures

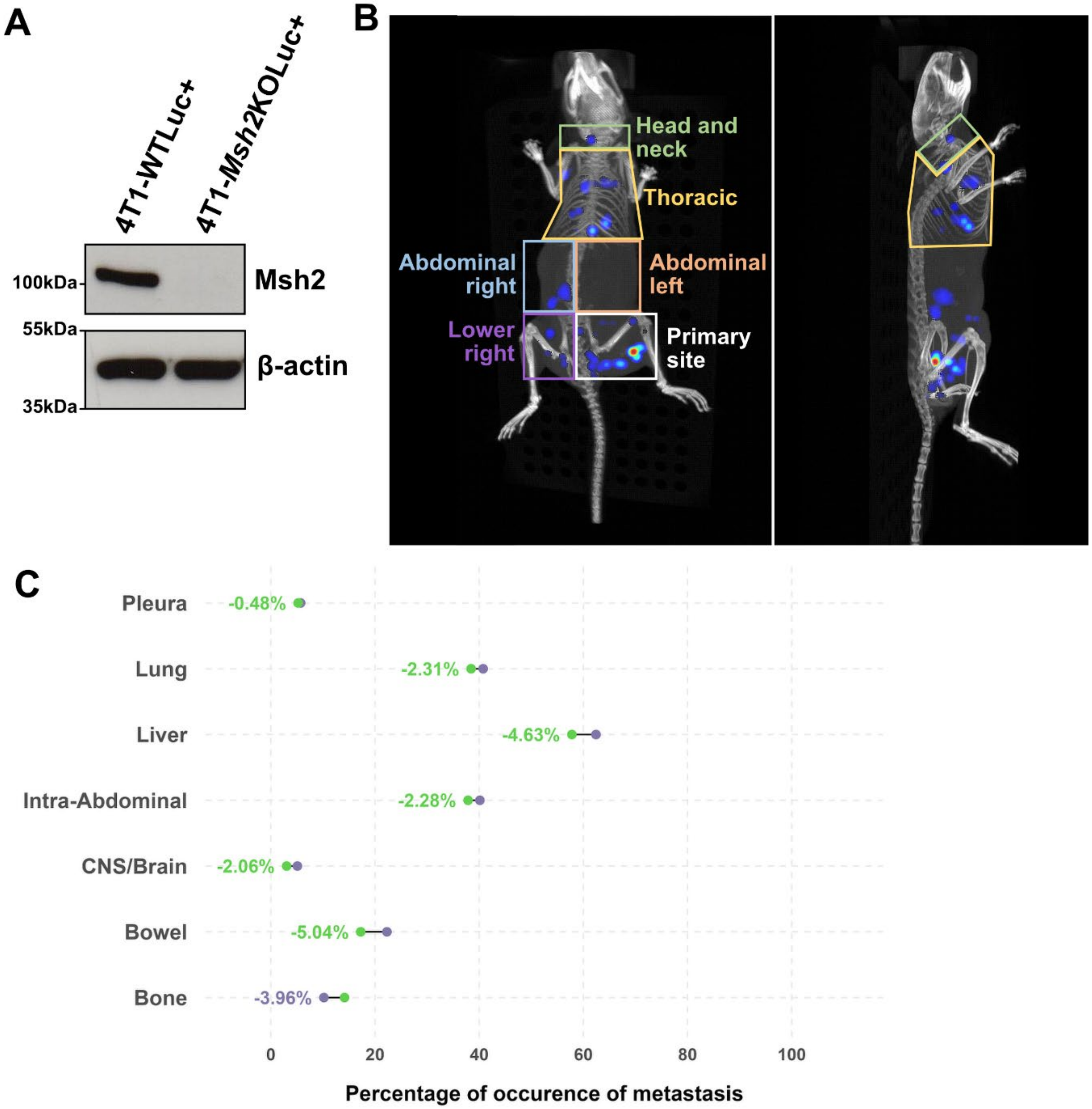

**Supplemental Figure S1.**

**A.** Validation of the loss of MSH2 protein detected by western blot in clone inactivated for *Msh2* gene by CRISPR/Cas9 in 4T1-Luc+ cell line and the parental 4T1-Luc+ cell line (WT). **B.** Schematic representation of the functional body sections used to localize metastasis-Luc+. **C.** Lollipop plot of the percentage of CRC MSS patient-bearing lesions at each organ (representative of the sites outlined in our cohort) according to TMB from the MSK-MET database. Only the MSS patients were retained for this analysis. Metastasis burden was transformed into metastasis occurrence (binary data), and the percentage of occurrence per organ was calculated for patients from the MSK-MET CRC database. Low TMB patients are displayed in green and High TMB patients are represented in purple.

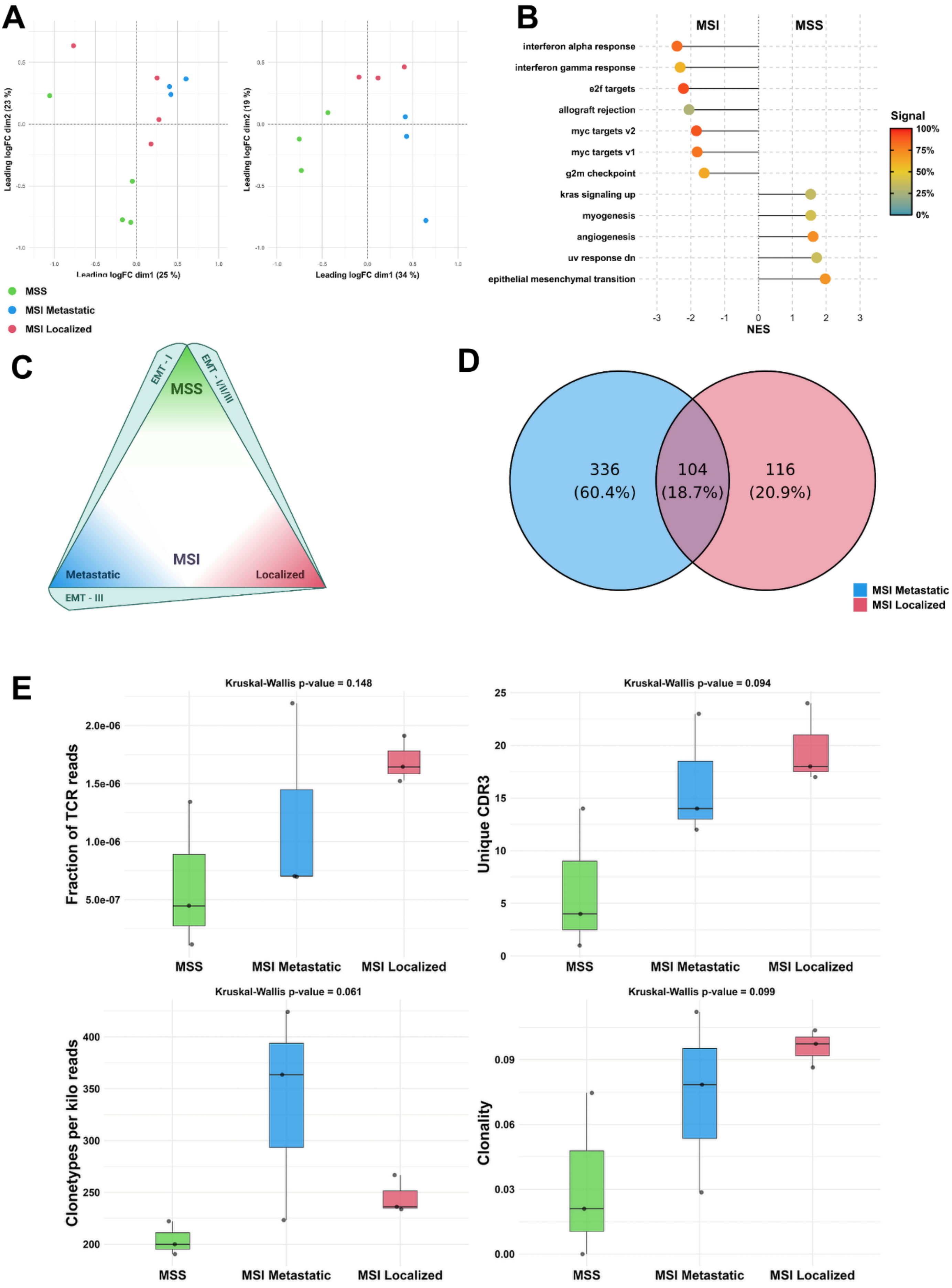

A
Leading logFC dim2 (23 %)
Leading logFC dim1 (25 %)
Leading logFC dim2 (19 %)
Leading logFC dim1 (34 %)
MSS
MSI Metastatic
MSI Localized
B
MSI
MSS
interferon alpha response
interferon gamma response
e2f targets
allograft rejection
myc targets v2
myc targets v1
g2m checkpoint
kras signaling up
myogenesis
angiogenesis
uv response dn
epithelial mesenchymal transition
NES
Signal
100%
75%
50%
25%
0%
C
EMT - I
EMT - I/II/III
MSS
MSI
Metastatic
Localized
EMT - III
D
336
(60.4%)
104
(18.7%)
116
(20.9%)
MSI Metastatic
MSI Localized
E
Kruskal-Wallis p-value = 0.148
Fraction of TCR reads
Kruskal-Wallis p-value = 0.094
Unique CDR3
Kruskal-Wallis p-value = 0.061
Clonetypes per kilo reads
Kruskal-Wallis p-value = 0.099
Clonality
MSS
MSI Metastatic
MSI Localized

**Supplemental Figure S2.**

**A.** PCoA plot of the RNA-seq samples before (left) and after (right) removal of two outlier samples. **B. GSEA differential enrichment of the Hallmark 50 gene sets.** Only significant gene sets are shown here. Positive NES signatures are enriched in MSS, and negative NES are enriched in MSI. Signal represents, in percentage, the number of genes present at the very beginning of the gene ranking made during GSEA. **C. Pairwise GSEA differential enrichment of the EMT signatures from the 41 cancer MPs.** Each group (MSS, MSI Metastatic and MSI Localized) is represented in each vertex. The signatures represented on the outline are enriched from thick to thin. EMT-I: "Full" mesenchymal programs compared to the other, that reflect partial EMT. EMT-II: Similar to the classical EMT program found in the hallmark 50 signatures (HALLMARK_EPITHELIAL_MESENCHYMAL_TRANSITION). EMT-III: A hybrid E-M program, including activation of mesenchymal and epithelial markers. Synonym of a plastic, aggressive state. **D.** Venn diagram of the private and shared neoantigens between the MSI Metastatic and MSI Localized groups. **E.** Boxplots of TCR repertoire metrics between MSS, MSI Metastatic and MSI Localized. PCoA: Principal Coordinate Analysis; EMT: Epithelial-Mesenchymal transition; NES: Normalized enrichment score.

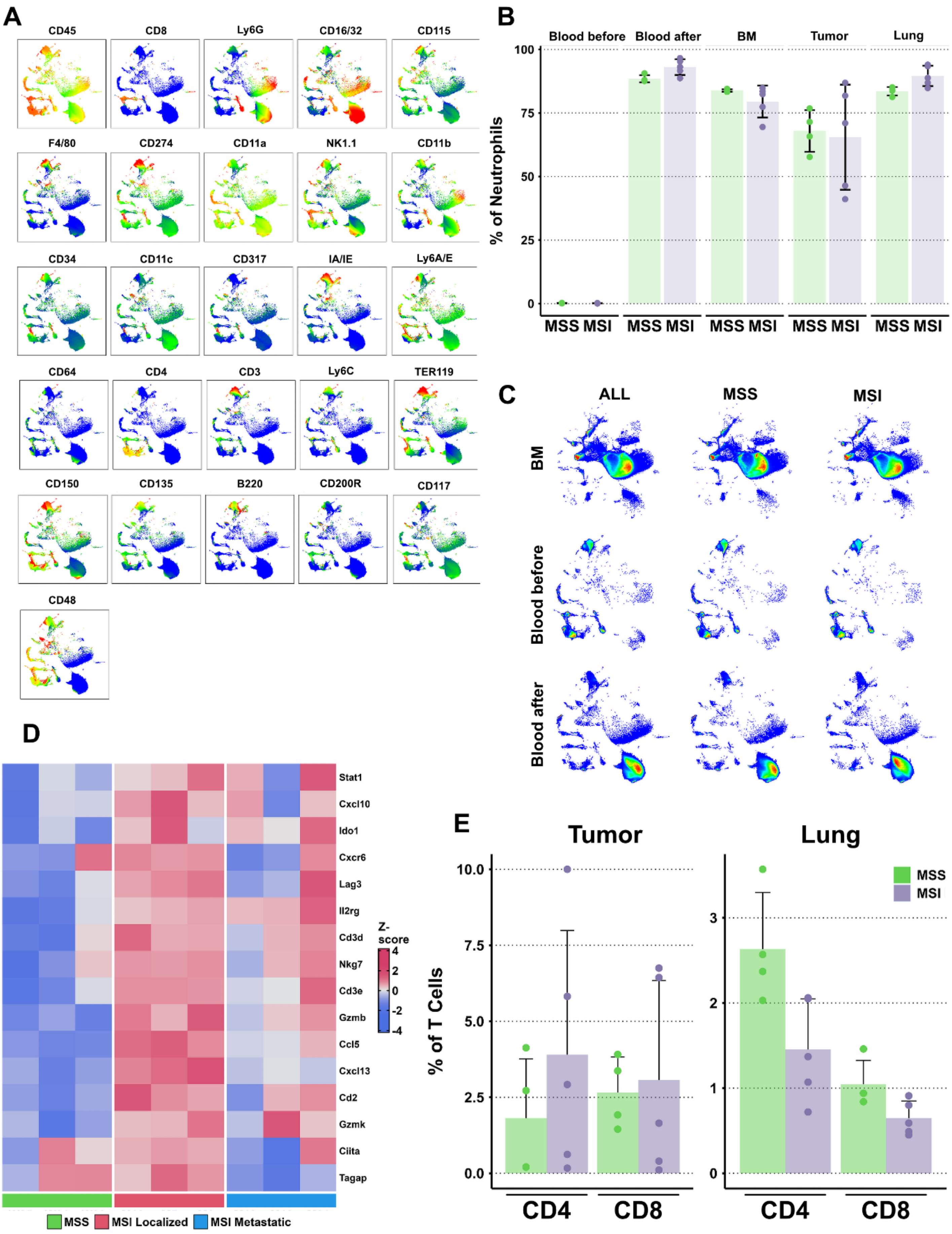
A
CD45
CD8
Ly6G
CD16/32
CD115
F4/80
CD274
CD11a
NK1.1
CD11b
CD34
CD11c
CD317
IA/IE
Ly6A/E
CD64
CD4
CD3
Ly6C
TER119
CD150
CD135
B220
CD200R
CD117
CD48
B
Blood before
Blood after
BM
Tumor
Lung
% of Neutrophils
100
75
50
25
0
MSS MSI
MSS MSI
MSS MSI
MSS MSI
MSS MSI
C
ALL
MSS
MSI
BM
Blood before
Blood after
D
Stat1
Cxcl10
Ido1
Cxcr6
Lag3
Il2rg
Cd3d
Nkg7
Cd3e
Gzmb
Ccl5
Cxcl13
Cd2
Gzmk
Ciita
Tagap
Z-score
4
2
0
-2
-4
MSS
MSI Localized
MSI Metastatic
E
Tumor
Lung
% of T Cells
10.0
7.5
5.0
2.5
0.0
3
2
1
0
MSS
MSI
CD4
CD8
CD4
CD8

**Supplemental Figure S3.**

**A.** UMAPs with color mapping representing the expression of each marker of the panel. Each panel is one marker with its associated expression on the UMAP. **B.** Neutrophils as percentage of all events in all immune compartments. **C.** UMAPs of events from the BM, blood before, blood after. "ALL" represents all samples combined. MSS and MSI represent the samples in their respective groups. **D.** Heatmap of the interferon-gamma signature genes expression from the RNA-seq experiment in Z-score-transformed of log2 count-per-million read of each gene. **E.** Bar graphs of T-CD4+ and T-CD8+ as percentages of CD45+ cells from MSS and MSI primary tumor and lung compartments. Error bars represent the standard deviation around the mean.
